# Aryl Hydrocarbon Receptor (Ahr) Pathway Drives TBBPA Induced Cartilage Development Defects

**DOI:** 10.64898/2026.09.07.749921

**Authors:** Kanchaka Senarath Pathirajage, Sunil Sharma, Tyler Johnson, Arnob Sarker, Subham Dasgupta

**Affiliations:** Department of Biological Sciences, Clemson University, Clemson, SC, USA; Environmental Toxicology Graduate Program, Clemson University, Clemson, SC, USA

## Abstract

Tetrabromobisphenol A (TBBPA), is one the most widely produced brominated flame retardant, detected in human matrices including cord plasma, raising concern over its impact on embryonic development. Our previous work demonstrated that TBBPA disrupts craniofacial cartilage development in zebrafish; however, inhibition of bone morphogenetic protein (BMP) signaling did not rescue these defects, suggesting the involvement of alternative molecular mechanisms. Transcriptomic profiling revealed significant upregulation of aryl hydrocarbon receptor (Ahr) signaling genes, including *cyp1a* and *cyp1c*, as well as reactive oxygen species (ROS) responsive genes, such as *nfe* and *gstp.* Consistent with these findings, KEGG pathway enrichment analysis identified significant enrichment of pathways involved in xenobiotic metabolism, cytochrome P450-mediated metabolism, and molecular docking predicted stronger binding of TBBPA to Ahr2 than the Ahr2 agonist TCDD. Then we investigated the role of Ahr signaling and ROS in TBBPA-induced craniofacial cartilage defects using microinjection of Ahr2 translation-blocking morpholino and the antioxidant N-acetylcysteine (NAC), respectively. Immunohistochemistry confirmed concentration-dependent induction of Cyp1a and ROS at environmentally relevant concentrations (≥0.005 µM). Ahr2 knockdown using a translation-blocking morpholino rescued TBBPA-induced alterations in Cyp1a expression, ROS levels, DNA damage, the expression of chondrogenesis-related markers (Sox10 and Sox2), and epithelial to mesenchymal transition (EMT) markers (E-cadherin, N-cadherin, and Snail2). Furthermore, Ahr2 knockdown rescued several TBBPA-induced alterations in craniofacial cartilage parameters. Co-exposure of TBBPA with the antioxidant NAC rescued the ROS and DNA damage and rescued only ceratohyal cartilage length, whereas other cartilage parameters remained disrupted. These findings establish a causal role for the Ahr-Cyp1a axis in TBBPA-induced craniofacial cartilage developmental toxicity and identify ROS as an important downstream contributor, indicating that both ROS-dependent and ROS-independent Ahr mechanisms underlined the observed TBBPA induced craniofacial defects.

## 1. Introduction

Tetrabromobisphenol A (TBBPA) is a widely used brominated flame retardant used in mainly electronics, household furniture, textiles, epoxy and polycarbonate resins for printed circuit boards due to their excellent fire-breaking feature that improve fire safety and reduce flammability ^1–4^. Because of high usage, global production volume of TBBPA and its derivatives is greater than 100,000 tons per year ^5^. Environmental monitoring and biomonitoring studies have detected TBBPA in indoor air, household dust, and human breast milk, indicating that inhalation, dust ingestion, and lactational transfer may represent important routes of human exposure ^6^. Furthermore, TBBPA has also been detected in food products, including meat, eggs, cheese, and fish, suggesting that dietary ingestion constitutes an additional pathway of human exposure ^6^. Consistent with these exposure pathways TBBPA levels has been traced in both adult plasma ^7^ and cord plasma ^5^. Therefore, these prenatal and early postnatal exposures underscore the need to examine the potential impacts of TBBPA on embryonic development, which has not yet been comprehensively investigated.

Leveraging zebrafish as a model, our previous work discovered that TBBPA targets histone acetylation during pre-pluripotent phases of development ^8^. Continuing this line of work, we focused on examining how TBBPA targets cell migration, dorsoventral patterning and organogenesis. We showed that TBBPA increased bone morphogenetic protein (BMP) signaling -a key regulator of dorsoventral patterning-in a concentration-dependent manner, as well as levels of several cell adhesion proteins and germ layer markers. ^9^. At later stages, TBBPA exposures induced concentration-dependent deficits in craniofacial cartilage development at 120 hours post fertilization (hpf) at low, environmentally relevant concentrations ^9^.

Cartilages are critical building block of body structures. Craniofacial cartilage development begins with the specification of cranial neural crest cells (CNCCs) ^9,10^. These CNCCs subsequently migrate into the pharyngeal arches, where they undergo mesenchymal condensation before differentiating into chondrocytes ^9,10^. Subsequently, chondrogenic transcription factors promote chondrocyte differentiation and cartilage matrix formation by activating extracellular matrix genes ^11^. Early embryonic development of the craniofacial skeleton, including the skull and jaw, is highly susceptible to environmental chemical exposure, which can disrupt the genetic regulation of chondrogenesis and lead to craniofacial and mandibular abnormalities. ^12^. In fact, the Centers for Disease Control and Prevention (CDC) reports that approximately 1 in every 33 infants born in the United States is affected by a congenital birth defect, a substantial proportion of which involve orofacial abnormalities.^13^. Therefore, examination of mechanisms of TBBPA-induced cartilage defects is crucial for understanding the full spectrum of effects of this ubiquitous flame retardant on developmental physiology. Along with cartilage defects, we also demonstrated that TBBPA exposure disrupted expression of key chondrogenic markers, including Sox2 and Sox10^14^. Additionally, although both phenotypic and protein-level evidence suggested BMP pathway overactivation, co-exposure with the BMP inhibitor dorsomorphin failed to rescue the defects suggesting involvement of alternative mechanisms ^9^ in driving craniofacial chondrogenesis defects.

The present study is aimed at examining these underlying mechanisms. To achieve this, we first conducted mRNA sequencing, which identified aryl hydrocarbon receptor (Ahr) signaling as a potential mediator of TBBPA toxicity. Following this, leveraging a reverse genetics approach, we investigated the role of Ahr pathway in driving chondrogenesis factors at an environmentally relevant concentration. Since Ahr induction is also associated with induction of reactive oxygen species, we also assessed how Ahr-induced ROS drives chondrogenesis defects. Collectively, this study fills a critical knowledge gap by establishing a causal role for Ahr signaling and ROS in TBBPA-induced craniofacial cartilage developmental toxicity.

## 2. Materials and Methods

### 2.1. Zebrafish rearing and embryo collection

Specific pathogen-free 5D wild type zebrafish were reared at a maximum density in non-chlorinated water glass tanks at a temperature of 27 ± 1 °C and a photoperiod of 14:10 h (light/dark) and were fed with artificial feed three times a day in Aquatic Animal Research Laboratory at Clemson University (AUP2022-0434 and AUP2023-0114). Embryos were transferred to petri dishes (∼30-50 embryos in a 100-mm Petri dish) containing E3 embryo medium.

### 2.2. Chemicals and morpholino

TBBPA (>99 % purity) (CAS #: 79-94-7), N-Acetyl-L-cysteine (>99 % purity) (CAS #: 616-91-1) and dimethyl sulfoxide (DMSO) were purchased from Sigma Aldrich.

### 2.3. TBBPA chemical exposure

Embryos were exposed at 6 h post-fertilization (hpf) to 0, 0.005, 0.05, 0.5 μM TBBPA in 10 mL glass beakers (*N* = 3 replicate beakers, 10 embryos per replicate). Each beaker contained 2 mL of TBBPA solution with 0.1 % DMSO as vehicle. Exposures were conducted until 120 hpf with daily renewal of the solutions. Based on our previous work ^9^, all exposure concentrations are within environmental relevance.

### 2.4. mRNA sequencing

To evaluate impacts on the mRNA expression, embryos (N=4, 20 embryos per replicate) were exposed to vehicle 0 or 5 μM TBBPA from 6 hpf to 24 hpf and incubated at 28 °C. At 24 hpf, 20 embryos per replicate pool (4 replicate pools) were collected into RINO 1.5 mL screwcap tubes, homogenized in RNAzol using a Bullet Blender Storm Pro (Next Advance, Inc., Troy, NY, USA) and initiated RNA extraction as described in Serradimigni et al., 2024. Sequencing was conducted on a Novaseq 600 (Illumina, San Diego, CA) (2 ×150 bp, ∼40 m reads/sample). Raw sequencing files will be deposited into NCBI Gene Expression Omnibus under accession # GSE310405.

### 2.5. Bioinformatic analysis of sequencing data

Raw sequencing reads in FASTQ format were processed using Novogene’s in-house Perl scripts. Adapter sequences, reads containing a high proportion of undetermined bases, and low-quality reads were removed to generate high-quality clean reads. Clean reads were aligned to the zebrafish reference genome (GRCz11) using HISAT2 v2.0.5. The number of reads mapped to each gene were quantified using feature counts v1.5.0-p3. Gene-expression levels will be reported as fragments per kilobase of transcript per million mapped reads (FPKM). Differential gene-expression analysis between control and TBBPA-exposed groups were performed using DESeq2 v1.20.0 with raw gene-count data. *P* values were adjusted for multiple comparisons using the Benjamini-Hochberg method to control the false-discovery rate. Genes with an adjusted *P* value below 0.05 were considered differentially expressed genes that are significantly altered by TBBPA exposure. Gene Ontology (GO) enrichment, biological and KEGG pathway analyses will be conducted using ShinyGO v0.85 to identify biological processes and molecular pathways significantly impacted by TBBPA exposure.

### 2.6. In silico molecular docking analysis

To investigate the interaction between Ahr2 and environmental toxicants, molecular docking analysis was performed between Ahr2 and TBBPA, along with 2,3,7,8-tetrachlorodibenzo-p-dioxin (TCDD)-a known Ahr2 agonist. The three-dimensional (3D) structure of Ahr2 was obtained from the PDB database ^15^. The 3D structures of TBBPA and TCDD were retrieved from the PubChem database ^16^. Prior to docking, receptor and ligand structures were prepared using standard procedures. Molecular docking was carried out using Auto Dock Vina ^17^ to predict the binding affinity and interaction patterns between Ahr2 and the selected ligands. The binding affinity scores were calculated in kcal/mol, where lower scores indicate stronger binding interactions.

### 2.7. Ahr2 translation blocking morpholino and microinjection

Morpholino antisense oligos were designed and manufactured by Gene Tools (Philomath, Oregon). A previously validated morpholino designed to inhibit Ahr2 protein translation (Ahr2 Morpholino: 5′-TGTACCGATACCCGCCGACATGGTT-3′) was used to reduce Ahr2 receptor expression (zfAhr2^18^). A standard control morpholino from Gene Tools (Control Morpholino: 5′-CCTCTTACCTCAGTTACAATTTATA-3′) served as the control. Both morpholinos were labeled with fluorescein at the 3′ end, enabling confirmation of successful injections using fluorescence microscopy. Morpholinos were diluted to 100 μM working solution in 1% DMSO. Approximately 3 nL of morpholino was injected into the yolk of embryos at the 1–2 cell stage (∼1 hpf) using a micro injector (Leica M80 Dissecting Microscope, Narishige micro manipulator, with BTX Microject 1000A injector). Embryos showing normal development and strong, uniform fluorescence at ∼ 4 hpf, confirming effective morpholino uptake, were selected and exposed to 2 mL of 0.5 μM of TBBPA exposure with 0.1 % DMSO as the vehicle up to the relevant developmental endpoint. (N = 5 replicate beakers, 10 embryos per replicate with 2.0 mL solution). At 24,72 or 120 hpf, morpholino injected embryos were phenotyped for developmental deformities. Subsets of embryos were allocated for ROS staining, and Alcian blue staining, whereas the remaining embryos were fixed overnight in 4% PFA, transferred to PBS and stored in 4 ^0^C for subsequent IHC.

### 2.8. Whole-mount immunohistochemistry (IHC)

Embryos were obtained from two experimental paradigms: (i) TBBPA concentration-response experiments, in which embryos were exposed to 0, 0.00005, 0.005, 0.5 or 5 μM TBBPA and (ii) Ahr2 morpholino experiments, in which control morphants and Ahr2 morphants were exposed to either 0.1% DMSO or 0.5 μM TBBPA and collected at 120 hpf. Subsequently embryos were fixed overnight in 4 % PFA and stored in PBS. Subsequently embryos were dechorionated and IHC was conducted according to prior protocols as explained in ^19^ with the following antibodies: anti-Cyp1a (mAbCRC4; 1:3), anti-E-cadherin (RR1; 1:2.5; Developmental Studies Hybridoma Bank/ DSHB), anti-N-cadherin (6B3;1:2.5, DSHB), anti-Snail2 (IE6; 1:2.5; DSHB), anti-Sox2 (1:100, Abcam), anti-Sox10 (1:200, Abcam) ,anti-Col2a1 (1:10, DSHB) and anti-phospho-H2a.x (1:100; Invitrogen-PA5-77995). After washing, embryos were counter stained in corresponding Alexa Fluor antibodies (1:500; Sigma) and visualized on an Echo Revolve Microscope (N = 10-15). Images were analyzed and quantified on Image J, with embryos stained with only secondary antibodies used as blanks.

### 2.9. Alcian blue cartilage staining for craniofacial cartilage development

Embryos were obtained from two experimental paradigms: (i) TBBPA concentration-response experiments, in which embryos were exposed to 0, 0.005, 0.05, or 0.5 μM TBBPA and (ii) Ahr2 morpholino experiments, in which control morphants and Ahr2 morphants were exposed to either 0.1% DMSO or 0.5 μM TBBPA, were collected at 120 hpf. Subsequently embryos were fixed in 4 % PFA for 2h at 4°C, followed by Alcian Blue staining (N = 10-12) to assess the impacts of TBBPA induced craniofacial cartilage development as described in Pathirajage et al., 2026 and Chen et al., 2026.

### 2.10. Reactive oxygen species (ROS) staining

Embryos were obtained from two experimental paradigms: (i) TBBPA concentration–response experiments, in which embryos were exposed to 0, 0.00005, 0.005, 0.5 or 5 μM TBBPA and (ii) Ahr2 morpholino experiments, in which control morphants and Ahr2 morphants were exposed to either 0.1% DMSO or 0.5 μM TBBPA, were collected at 24 hpf. Subsequently embryos were incubated for 20 min with CM-H₂DCFDA (20 µM; Abcam) stain to assess ROS production induced by TBBPA exposure at 24 hpf as described in Ferdous et al., 2026.After staining, embryos were washed three times with PBS buffer and then transferred to embryo media followed by imaging using an Echo Revolve Microscope (N = 10-15). Images were analyzed and quantified on Image J.

### 2.11. TBBPA - N-acetylcysteine (NAC) co exposure

At 6 hpf, embryos were co-exposed to TBBPA (0.5 µM) and NAC (100 µM) ^22^. Staining for ROS, DNA damage and craniofacial deficits were conducted as described previously.

### 2.12. Statistics

All statistical estimations were carried out in GraphPad Prism 9. Statistical significance was calculated using one-way ANOVA, followed by Dunnett’s post hoc tests for comparison of each treatment concentration to the DMSO controls (*p* < 0.05). Detailed results from statistical tests (ANOVA *p* values and Dunnett’s p values) are included within each figure. For co-exposure experiments with Ahr2 morpholino or NAC, a 2-way ANOVA, followed by Sidak’s test was used to determine the statistically significant differences (*p* < 0.05).

## 3. Results

### 3.1. TBBPA decreases craniofacial cartilage development

Following up on our previous work ^9^, we conducted Alcian blue staining at 120 hpf, with an expanded toolset of morphometric parameters. Our data revealed that TBBPA exposure significantly reduced lower jaw length (LJL), ceratohyal cartilage length (CCL), and total craniofacial cartilage area at both 0.05 and 0.5 μM and inter cranial distance only at 0.5 μM **(Fig. 1A-D)**. TBBPA also altered all measured cartilage angles, including the PQ-Meckel’s angle, the CH angle (ceratohyal), and the CH-PQ angle, while Meckel’s angle remained unchanged **(Fig. 1E-H).** Notably, the ceratohyal angle was the most sensitive, showing an increase even at the lowest TBBPA concentration (0.005 μM), whereas the CH-PQ and PQ-M angles increased at 0.05 and 0.5 μM, respectively **(Fig. 1E-H)**.

**Fig. 1.**
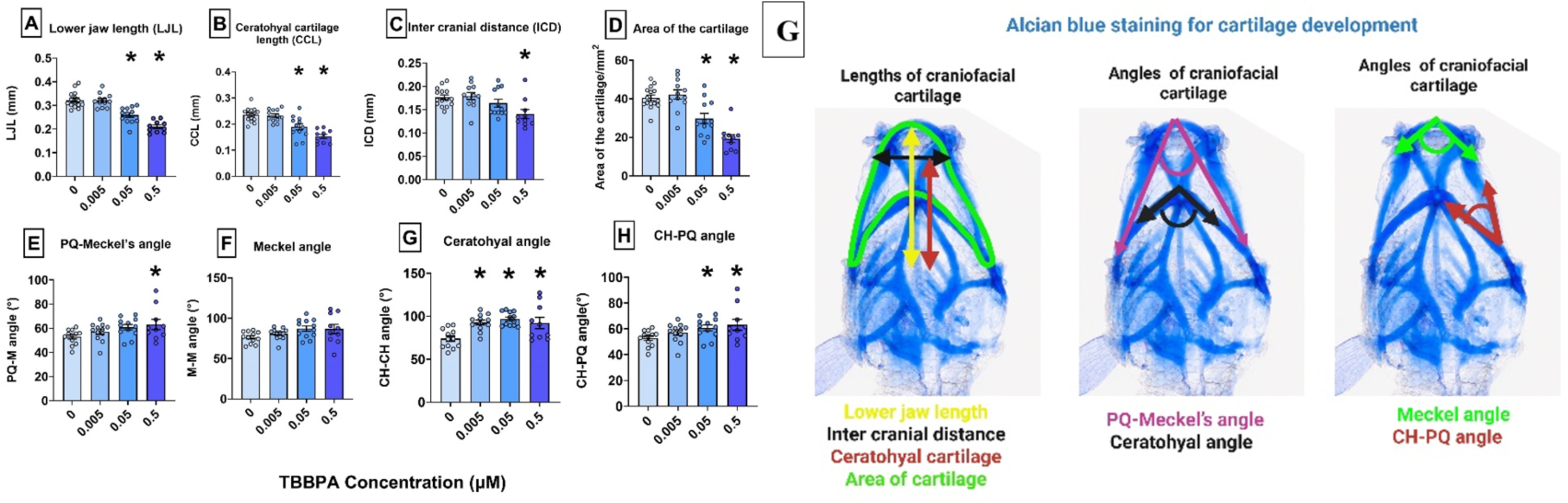
TBBPA decreased craniofacial cartilage development at 120 hpf. (**A)** Lower jaw length (LJL). **(B)** Intercranial distance (ICD). **(C)** Ceratohyal cartilage length (CCL). **(D)** Area of cartilage. (**E)** PQ-Meckel’s angle (PQ-M angle). **(F)** Meckel angle (M-M). **(G)** Ceratohyal angle (CH-CH). **(H)** CH-PQ Angle. **(G)** Length and Angle cartilage parameters. Asterisk (*) denotes statistically different (*p* < 0.05) from 0 μM based on Dunnett’s post hoc test following 1-way ANOVA.

### 3.2. TBBPA induces differential gene expression and significantly affects numerous biological pathways

Volcano plot analysis revealed that TBBPA exposure induced differential gene expression, with a total of 2,590 differentially expressed genes (DEGs) were identified, including 1,222 downregulated genes and 1,368 upregulated genes **(Fig. 2A)**. Functional enrichment analysis **(Fig. 2B)** indicated that TBBPA exposure initially activated xenobiotic-metabolizing and cellular stress-response pathways, as demonstrated by the significant enrichment of xenobiotic metabolism by cytochrome P450 and p53 signaling. In parallel, enrichment of p53 pathway suggests TBBPA-induced apoptosis-mediated developmental toxicity.

**Fig. 2A.**
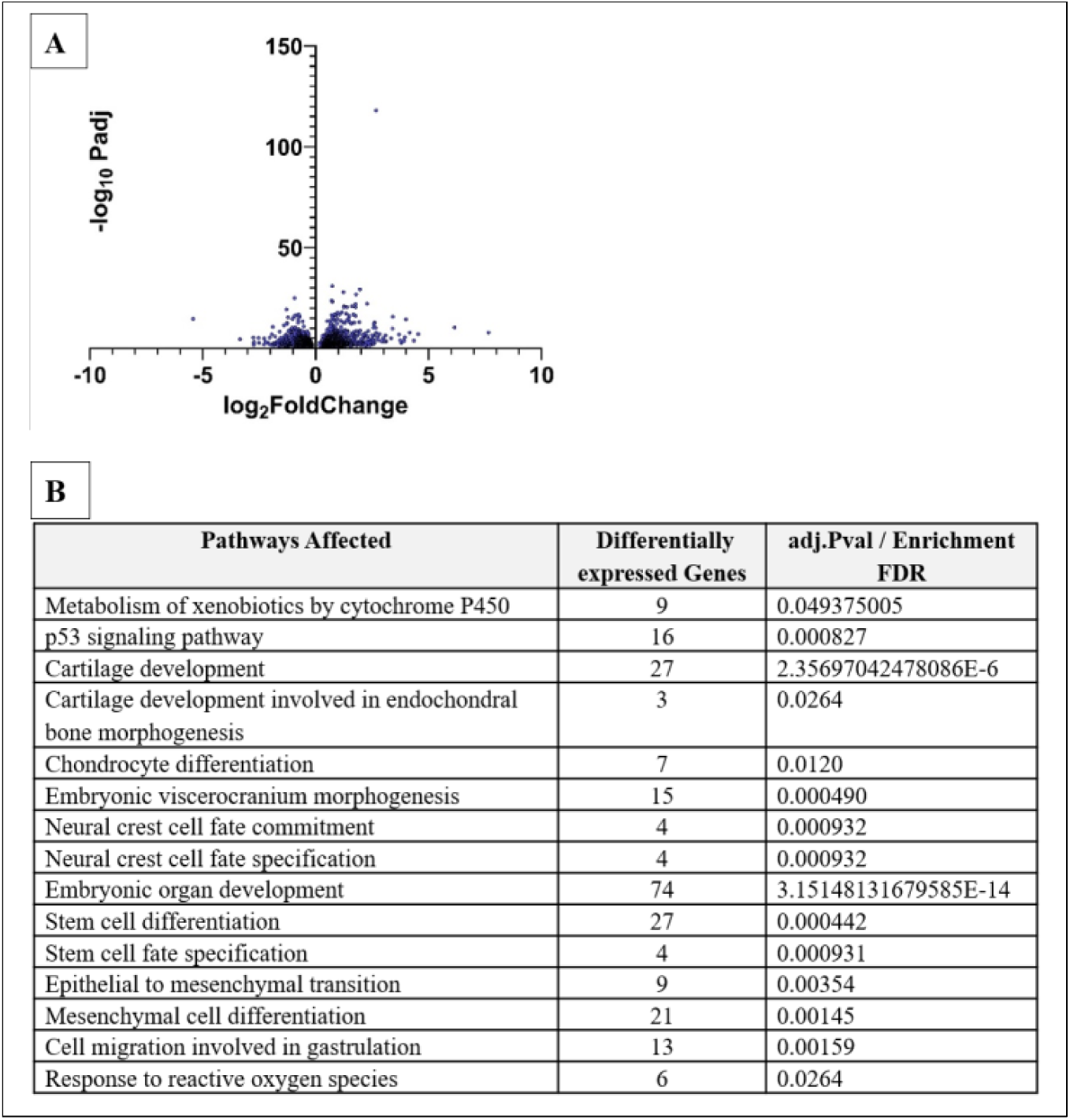
Volcano plot of TBBPA-induced differential gene expression at 24 hpf. Fig. 2B. Pathways affected according to KEGG and biological pathway analysis at 24 hpf.

### 3.3. TBBPA induces gene expression associated with Ahr signaling, cartilage and oxidative stress

Transcriptomic profiling by mRNA sequencing from 6 to 24 hpf **(Table 1)** revealed that TBBPA exposure significantly increases Ahr signaling. Several Ahr responsive genes were strongly upregulated, including *cyp1a* (log₂ fold change = 6.16), *cyp1c1* (log₂ fold change = 3.74), and *ahrra* (log₂ fold change = 2.32), confirming that TBBPA induces Ahr gene expression. TBBPA exposure also increased the expression of cartilage-associated genes, including *mmp9* (log₂ fold change = 3.09), *foxqi1* (log₂ fold change = 2.59), and *slincR* (log₂ fold change = 2.09). In addition, several genes involved in reactive-oxygen-species responses were upregulated, including *nfe* (log₂ fold change = 1.76), *gstp* (log₂ fold change = 1.79), and *noxo1a* (log₂ fold change = 1.21).

**Table 1.**
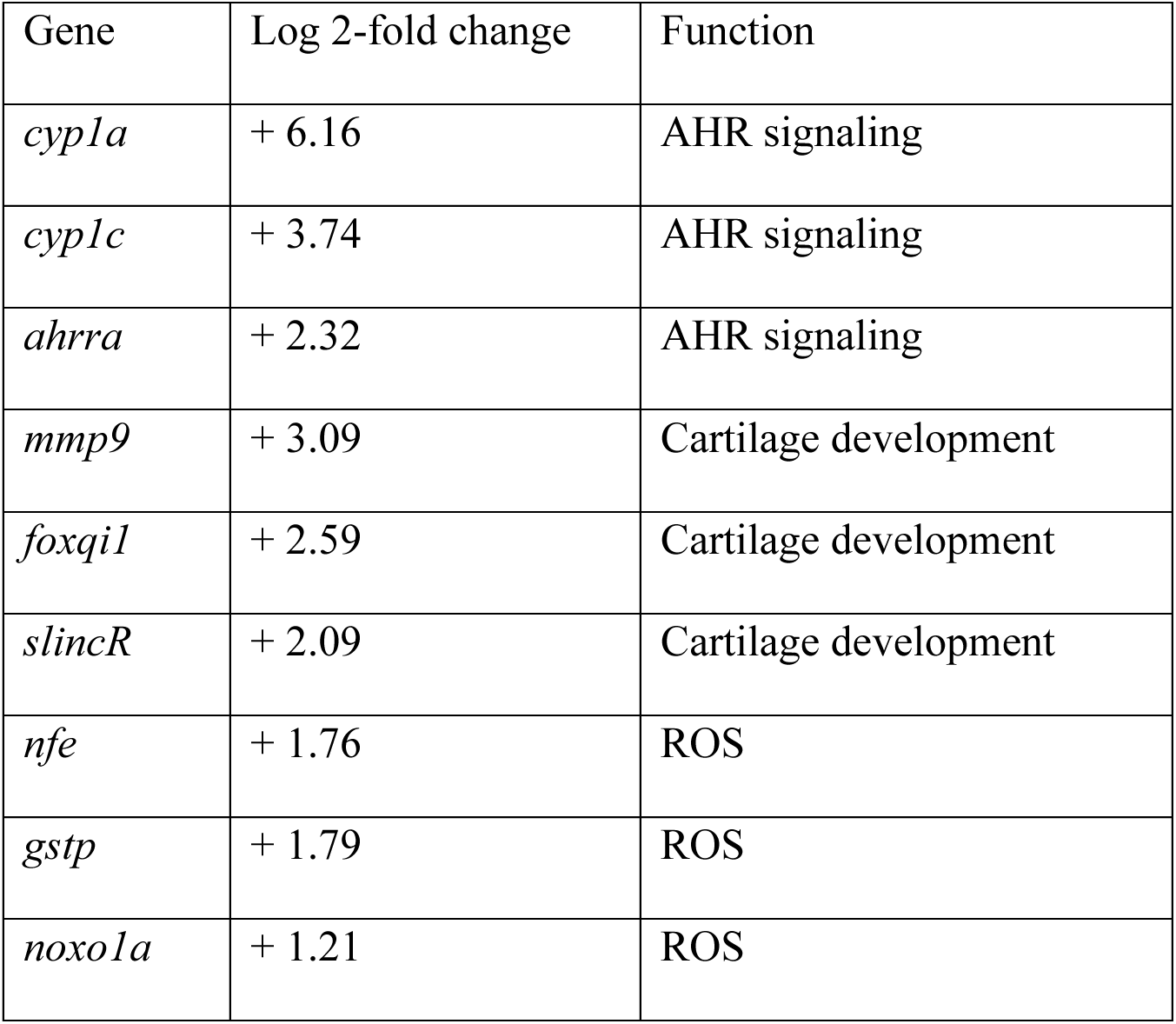
mRNA seq. TBBPA altered the gene expression of AHR signaling cartilage development and ROS.

### 3.4. Molecular docking analysis of Ahr2 with TBBPA and TCDD

To investigate the binding interactions between environmental toxicants and Ahr, molecular docking analysis was performed using TBBPA and TCDD as ligands. The docking results demonstrated favorable binding affinities for both compounds toward Ahr2. Among the two ligands, TBBPA exhibited the strongest interaction with Ahr2, showing a binding affinity of -7.5 kcal/mol, whereas TCDD displayed a binding affinity of -6.6 kcal/mol. The lower binding energy of the Ahr2-TBBPA complex suggests a more stable interaction compared with the Ahr2-TCDD complex. These findings indicate that both TBBPA and TCDD can effectively bind to Ahr2, suggesting a potential role for Ahr-mediated effects on zebrafish development.

### 3.5. TBBPA exposure significantly increased Cyp1a expression and ROS 24 and 72 hpf

Consistent with mRNA-sequencing results TBBPA exposure produced a concentration-dependent increase in Cyp1a protein expression **(Fig. 3)**. At 24 hpf, Cyp1a protein levels were significantly elevated at lower concentration down to 0.005 µM TBBPA **(Fig. 3A)** whereas it was elevated at even 0.00005 µM TBBPA at 72 hpf **(Fig. 3B).** Additionally, CH_2_DCFDA staining at 24 hpf showed that TBBPA increased ROS starting at 0.005 µM in a concentration dependent manner **(Fig. 4)**.

**Fig. 3.**
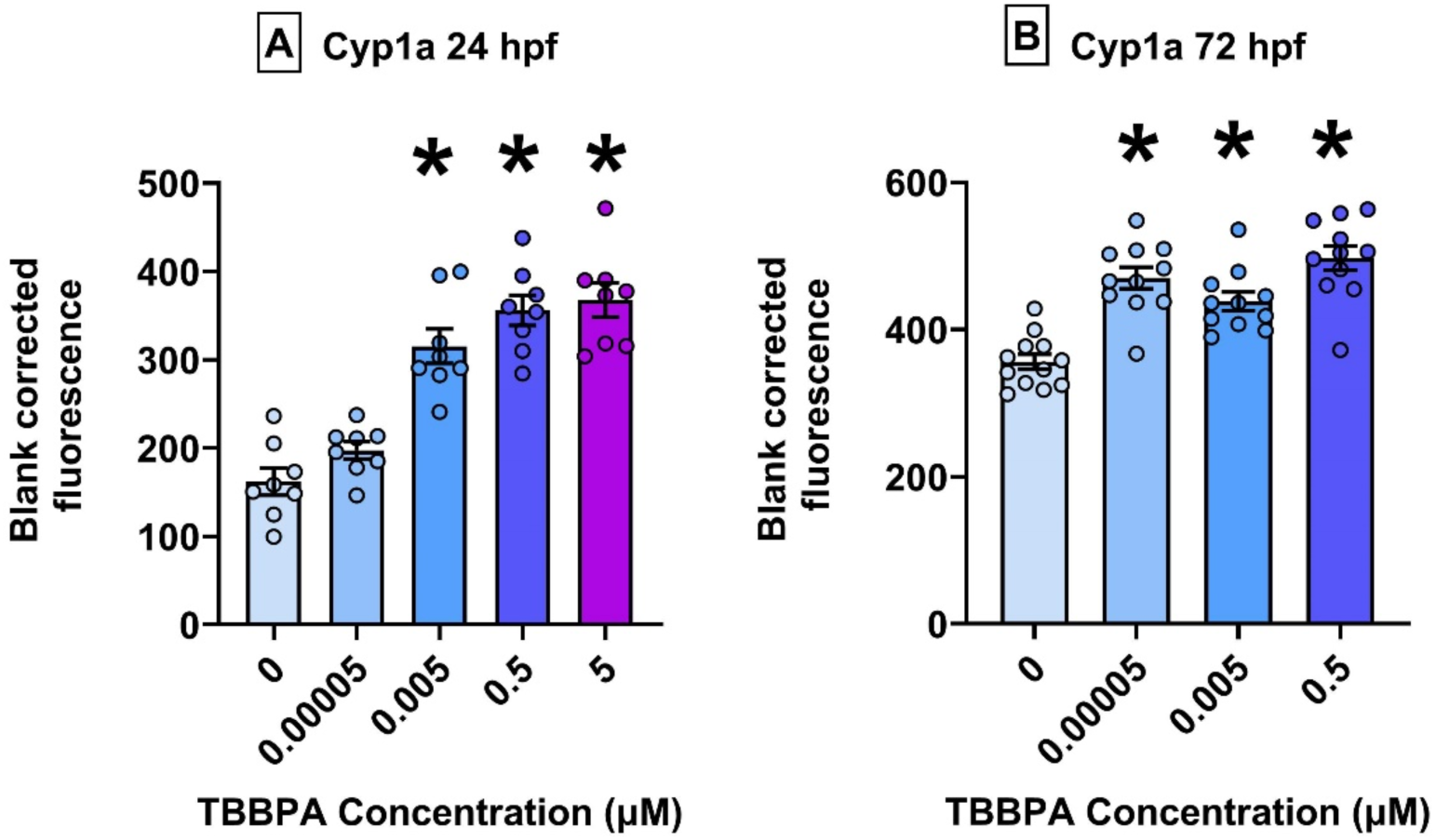
TBBPA increased Cyp1a expression in a concentration-dependent manner. **(A)** 6–24 hpf exposure **(B)** 6–72 hpf exposure. Asterisk (*) denotes statistically different (*p* < 0.05) from 0 μM based on Dunnett’s post hoc test following 1-way ANOVA.

**Fig. 4.**
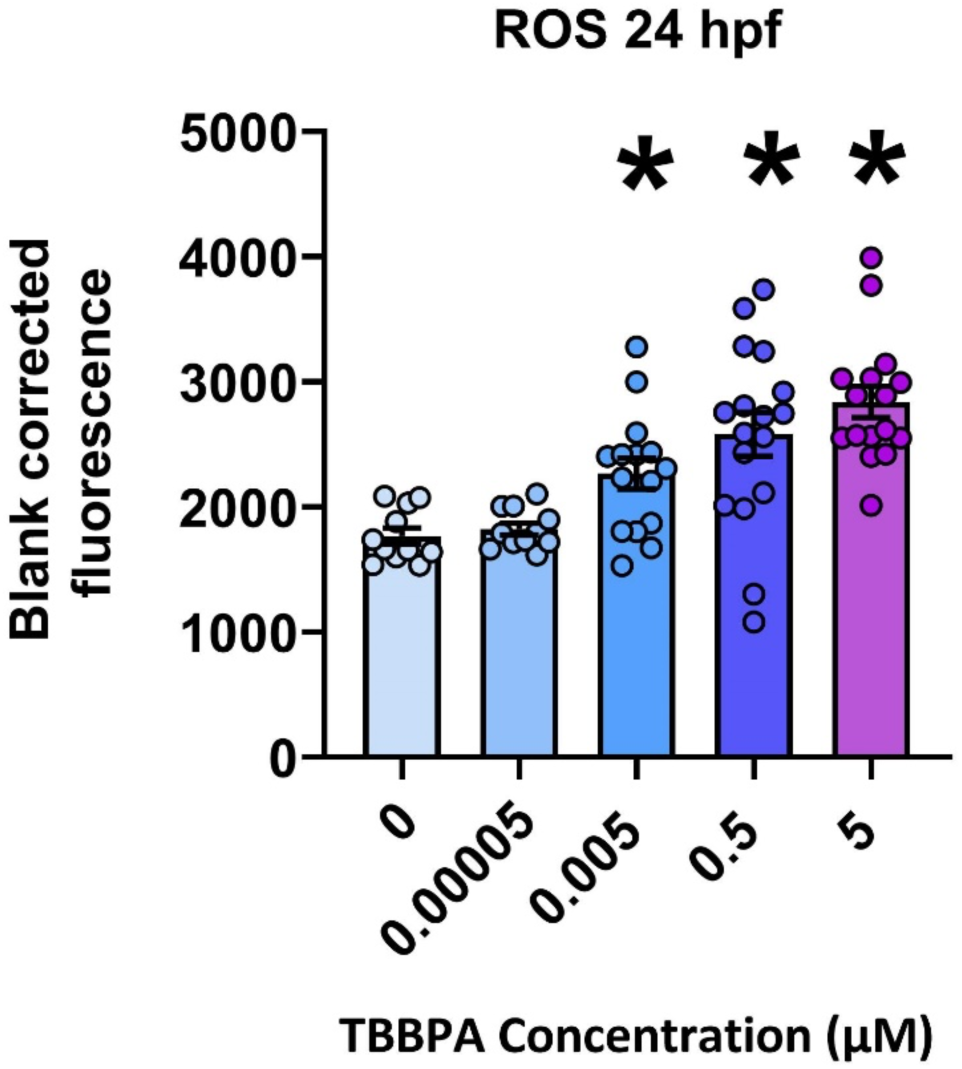
TBBPA increased ROS in a concentration-dependent manner 6-24 hpf. Asterisk (*) denotes statistically different (*p* < 0.05) from 0 μM based on Dunnett’s post hoc test following 1-way ANOVA.

### 3.6. Ahr2 knockdown rescued TBBPA-induced upregulation of Cyp1a expression at 24 and 72 hpf

Based on IHC at 24 and 72 hpf, exposure to 0.5 µM TBBPA significantly rescued the TBBPA-induced increases in Cyp1a expression in Ahr2 morphants compared to control morphants **(Fig. 5)**.

**Fig. 5.**
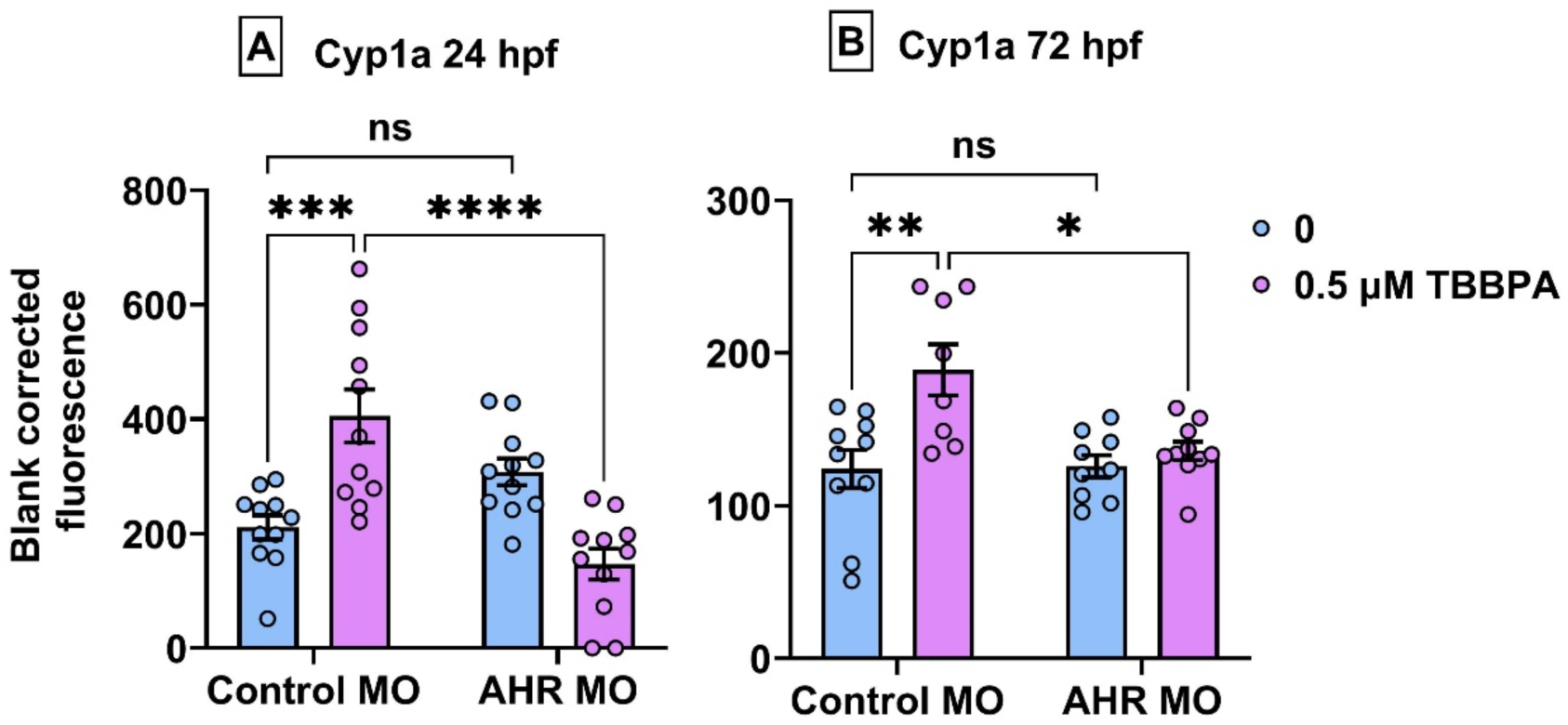
AHR2 morphants rescued the TBBPA indued increase Cyp1a expression (A) 6–24 hpf exposure (B) 6–72 hpf exposure. Asterisk (*) denotes statistically different (*p* < 0.05) from treatments based on Sidak post hoc test following 2-way ANOVA.

### 3.7. Ahr2 knockdown rescued TBBPA-induced craniofacial cartilage development at 120 hpf

Based on Alcian Blue staining at 120 hpf, Ahr2 knockdown significantly rescued TBBPA-induced disruptions in ceratohyal cartilage length and lower jaw length in Ahr morphants, but not area of cartilage or intercranial distance **(Fig. 6 A-D)**. Similarly, Ahr2 knockdown significantly rescued TBBPA-induced disruptions in the PQ-Meckel’s angle and the CH angle, but not Meckel’s angle and the CH-PQ angle **(Fig. 6E-H).**

**Fig. 6.**
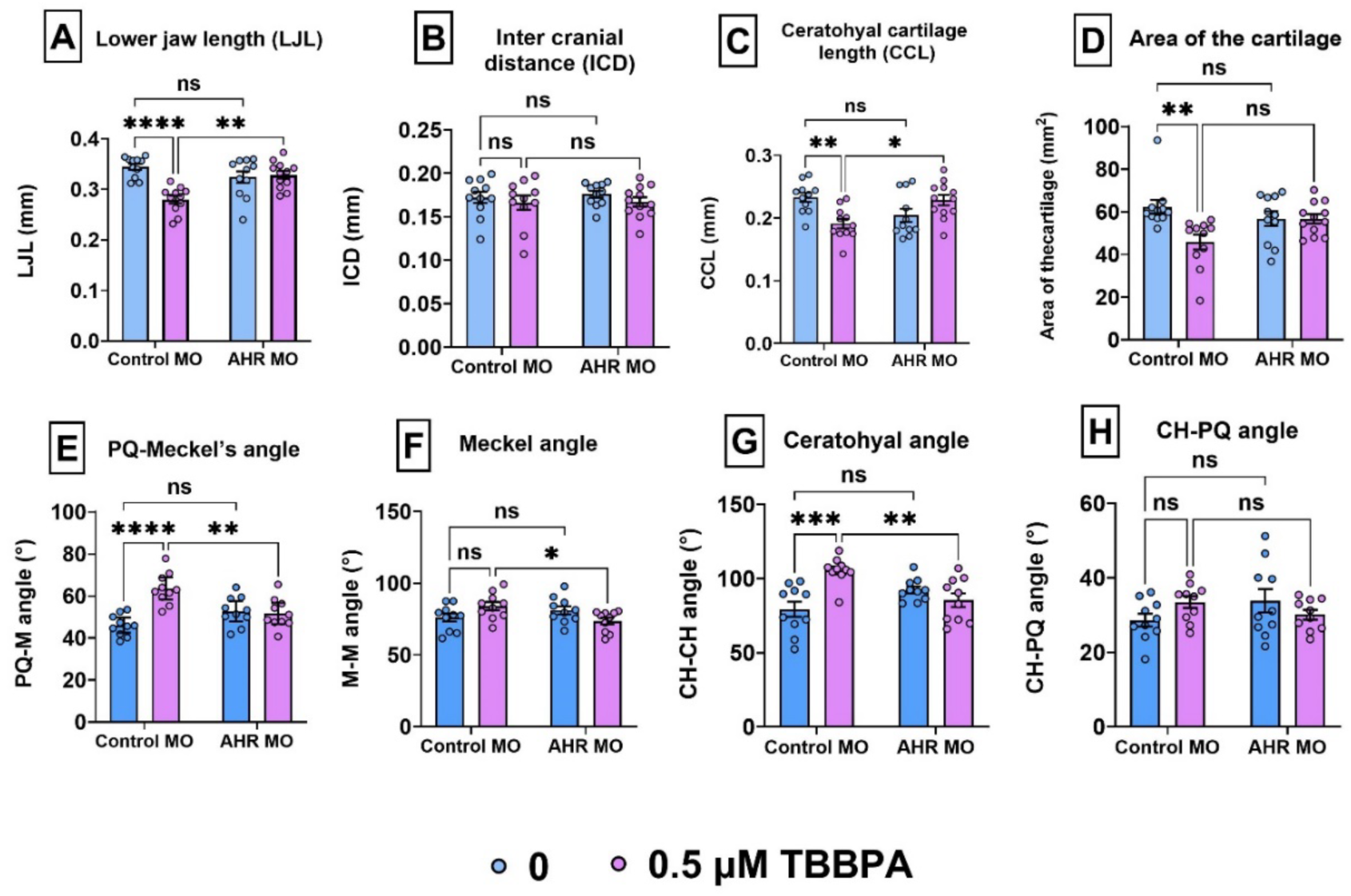
AHR2 morphants rescued the TBBPA indued cartilage disruption at 120 hpf. (**A)** Lower jaw length (LJL). **(B)** Intercranial distance (ICD). **(C)** Ceratohyal cartilage length (CCL). **(D)** Area of cartilage. (**E)** PQ-Meckel’s angle (PQ-M angle). **(F)** Meckel angle (M-M). **(G)** Ceratohyal angle (CH-CH). **(H)** CH-PQ Angle. Asterisk (*) denotes statistically different (*p* < 0.05) from treatments based on Sidak post hoc test following 2-way ANOVA.

### 3.8. Ahr2 knockdown rescued the TBBPA-induced alteration of markers in epithelial-to-mesenchymal transition (EMT) and cartilage development at 24 and 72 hpf

Based on IHC, Ahr2 knockdown significantly rescued the TBBPA-induced disruptions in EMT markers E-cadherin in Ahr2 morphants, whereas a rescue was not observed for N-cadherin and Snail2 **(Fig. 7A-C)**. Ahr2 knockdown also significantly rescued the TBBPA-induced disruptions in chondrogenesis markers such as Sox2, Sox10 at 24 hpf and Col2a1 at 72 hpf in Ahr2 morphants compared to control morphants **(Fig. 7D-F)**.

**Fig. 7.**
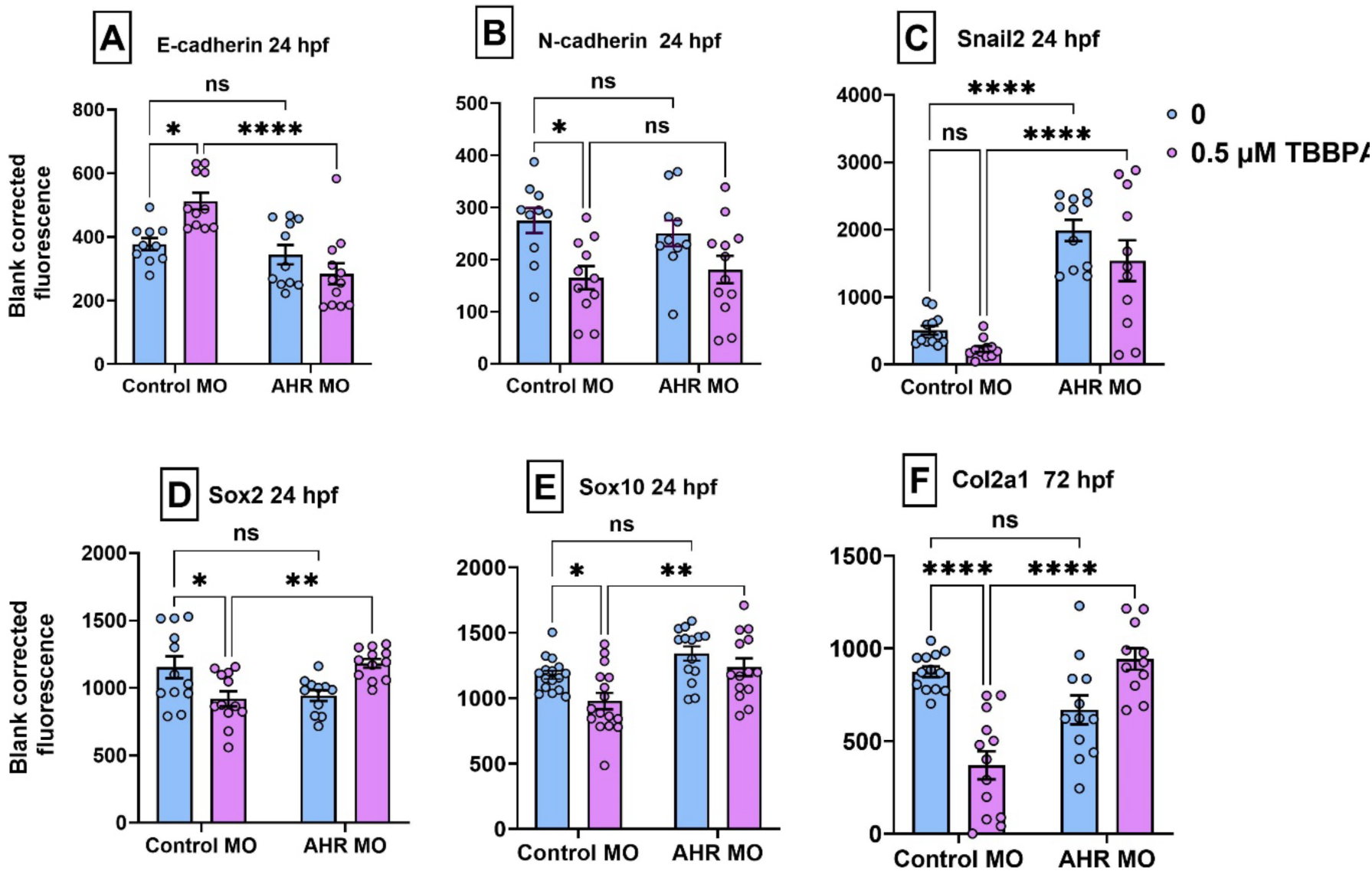
AHR2 morphants rescued the TBBPA induced alteration of markers of craniofacial cartilage development and EMT factors at 24 hpf and 72 hpf. **(A)** Sox2. **(B)** Sox10. **(C)** E-cadherin. **(D)** N-cadherin. **(E)** Snail2. Asterisk (*) denotes statistically different (p < 0.05) from treatments based on Sidak post hoc test following 2-way ANOVA

### 3.9. Ahr2 knockdown rescued TBBPA-induced increase in ROS and DNA damage at 24 hpf

Based on CH_2_DCFDA staining and IHC, exposure to Ahr2 knockdown significantly rescued both TBBPA-induced increase of ROS and DNA damage in Ahr2 morphants compared to control morphants **(Fig. 8A-B)**.

**Fig. 8.**
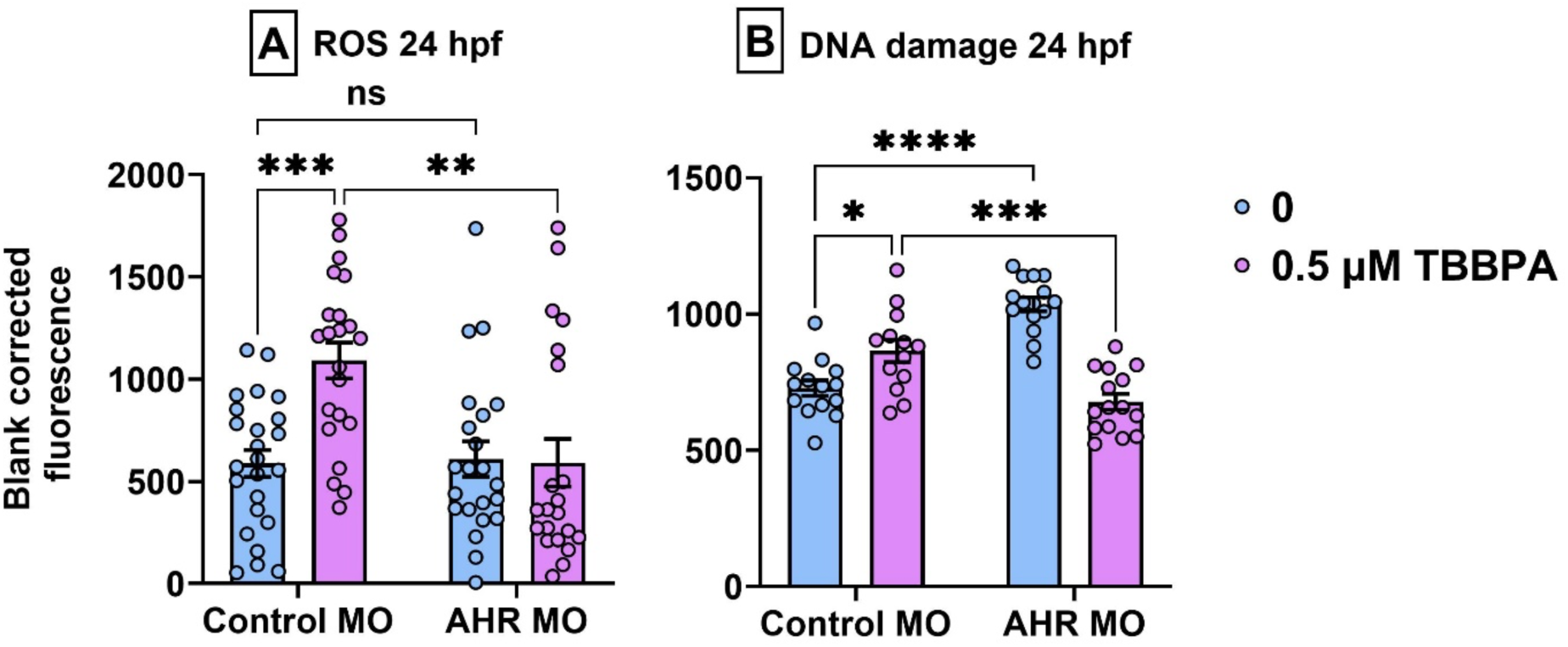
AHR2 morphants rescued the TBBPA induced ROS and DNA damage at 24 hpf. Asterisk (*) denotes statistically different (*p* < 0.05) from treatments based on Sidak post hoc test following 2-way ANOVA.

### 3.10. NAC rescued the TBBPA-induced increase in ROS and altered DNA damage at 24 hpf

Based on CH_2_DCFDA staining at 24 hpf and IHC on DNA damage, exposure to NAC significantly rescued TBBPA-induced increase of ROS and altered DNA damage in NAC exposed embryos compared to NAC unexposed **(Fig. 9A-B)**.

**Fig. 9.**
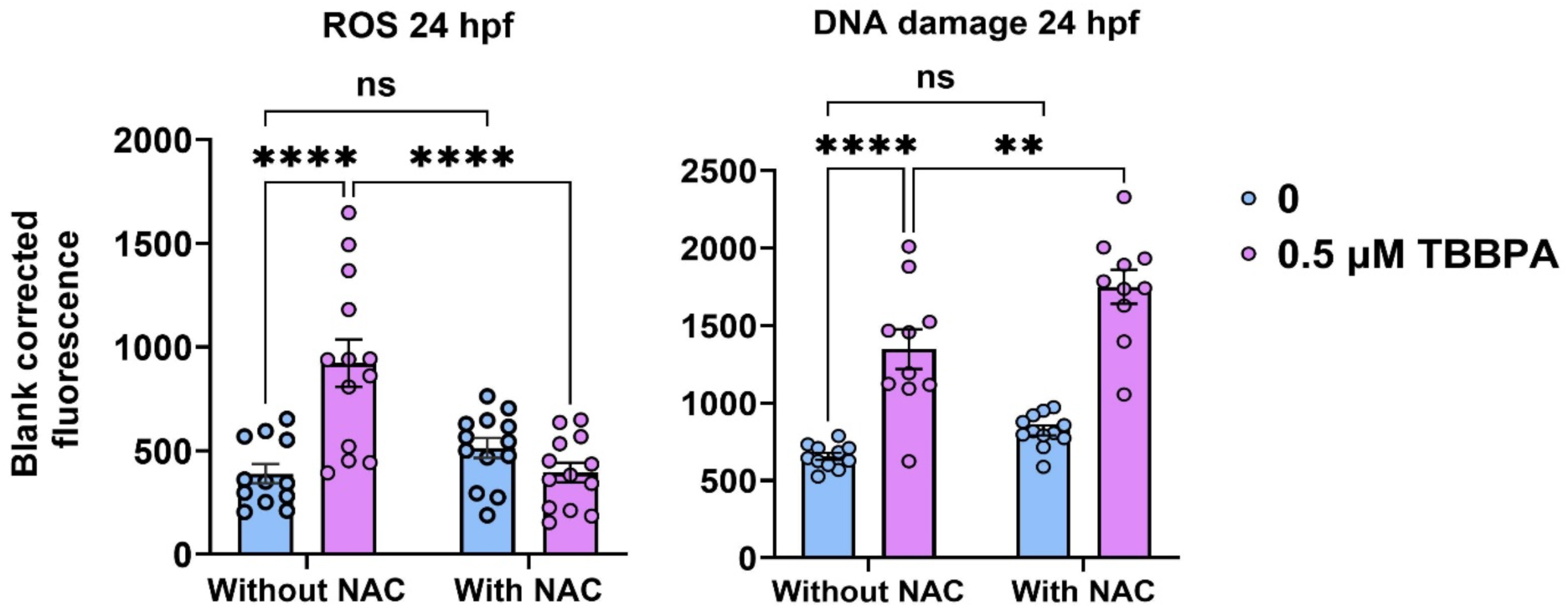
NAC rescued the TBBPA induced ROS and altered DNA damage at 24 hpf. Asterisk (*) denotes statistically different (*p* < 0.05) from treatments based on Sidak post hoc test following 2-way ANOVA.

### 3.11. NAC rescued the TBBPA-induced craniofacial development at 120 hpf

Based on Alcian Blue staining at 120 hpf, Ahr2 knockdown significantly rescued the TBBPA-induced disruptions in ceratohyal cartilage length in NAC exposed embryos. Interestingly the lower jaw length, intercranial distance and area of the cartilage failed to be rescued in NAC co-exposures **(Fig. 10A-D)**. As with cartilage length, NAC rescued coexposures only TBBPA-induced disruption in the PQ-Meckel’s angle. Notably, this angle was also specifically increased by NAC exposure alone, suggesting that it is particularly sensitive to NAC. In contrast, the remaining cartilage angles were not rescued in NAC-exposed embryos compared with NAC-unexposed embryos, including the Meckel’s angle, CH angle, and CH-PQ angle **(Fig. 10E-H)**.

**Fig. 10.**
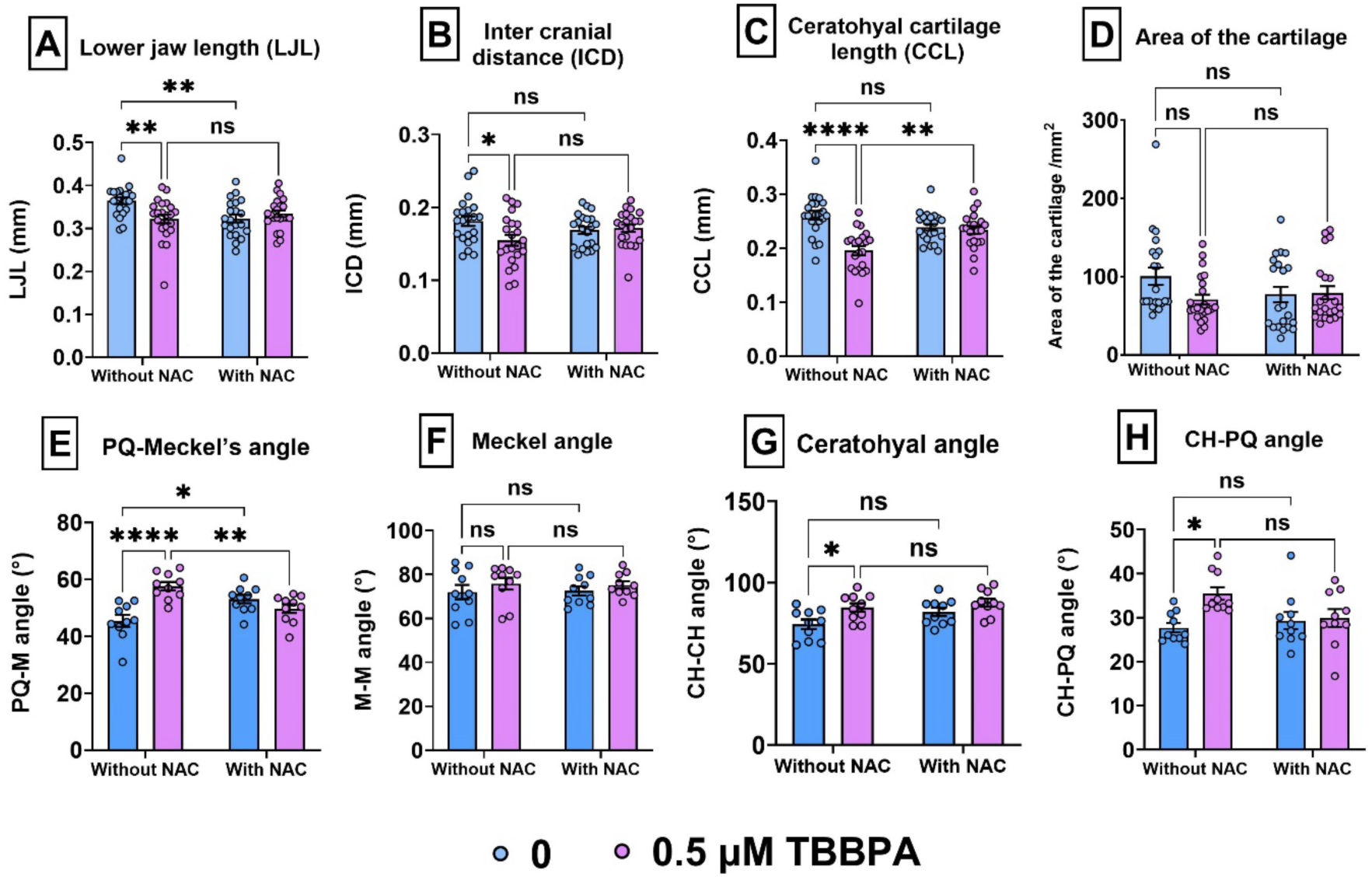
NAC rescued the TBBPA induced cartilage disruption at 120 hpf. (**A)** Lower jaw length (LJL). **(B)** Intercranial distance (ICD). **(C)** Ceratohyal cartilage length (CCL). **(D)** Area of cartilage. (**E)** PQ-Meckel’s angle (PQ-M angle). **(F)** Meckel angle (M-M). **(G)** Ceratohyal angle (CH-CH). **(H)** CH-PQ Angle. Asterisk (*) denotes statistically different (*p* < 0.05) from treatments based on Sidak post hoc test following 2-way ANOVA.

## 4. Discussion

Our previous study demonstrated that TBBPA disrupts bone morphogenetic protein (BMP) signaling and epithelial-to-mesenchymal transition (EMT), resulting an impaired germ layer formation ^19^. These effects were particularly evident in neural crest cell (NCC)-derived ectodermal development and subsequent ectoderm-derived cartilage formation, both of which were adversely affected by environmentally relevant TBBPA concentration. The environmental relevance of this exposure range is supported by our previous LC-MS analysis, which showed that a nominal 5 μM TBBPA exposure resulted in an embryonic tissue burden of 62 ± 41 ng/g ^19^, comparable to levels reported in human serum and umbilical cord plasma ^4^. Together, these findings indicate that environmentally realistic TBBPA exposures may adversely affect NCC-dependent craniofacial cartilage development.

Notably, BMP inhibition failed to rescue the cartilage abnormalities caused by TBBPA exposure, suggesting that mechanisms other than BMP signaling may be involved in TBBPA-induced cartilage effects. Therefore, we performed mRNA sequencing to identify additional molecular pathways potentially responsible for TBBPA-induced disruption of cartilage development. mRNA-seq data **(Table 1)**, coupled with whole mount IHC of Cyp1a **(Fig. 3)**, showed that TBBPA significantly upregulated the aryl hydrocarbon receptor (Ahr) signaling pathway, which regulates xenobiotic metabolism and mediates responses to environmental contaminants such as dioxins and PAHs ^23^. These Ahr-related findings are consistent with the broader KEGG and biological pathway enrichment results **(Fig. 2B)** , which suggest that TBBPA affects both developmental and xenobiotic-response pathways ^19^. The enrichment of drug metabolism and metabolism of xenobiotics by cytochrome P450 pathways suggest activation of Ahr signaling mechanisms. Consistent with that, our molecular docking studies showed that TBBPA binds to Ahr2. We therefore hypothesized that Ahr activation may act as an upstream mechanism driving TBBPA-induced cartilage defects. Prior studies have shown that a potent Ahr ligand, TCDD impairs normal jaw cartilage development via a Sox9b-dependent pathway ^24^ and disrupt cranial NCC-derived craniofacial bone development via Ahr/ERα signaling crosstalk, leading to teratogenic outcomes ^25^ indicating that activation of Ahr signaling can disrupt cartilage development and craniofacial morphogenesis.

One major consequence of Ahr signaling activation is the generation of excess reactive oxygen species (ROS), including superoxide and hydroxyl radicals ^26^. Excessive ROS production can lead to irreversible oxidative damage to DNA, RNA, proteins, lipids, and carbohydrates, ultimately triggering apoptosis or necrosis ^27^ ^23,28,29^. Activation of Ahr-dependent detoxification mechanisms by TCDD, PAHs, PCBs, and UV radiation has been strongly associated with increased reactive ROS production, oxidative stress and DNA damage ^23,30–32^. Indeed, upregulation of ROS and p53 genes **(Table 1)**, increased ROS and DNA damage levels detected by H_2_DCFDA staining **(Fig. 4, 8)** and the reversal of these levels by Ahr2 knockdown demonstrate that TBBPA-induced activation of the Ahr2-Cyp1a axis leads excess ROS and DNA damage. These, in turn, can have downstream implications on developmental pathways, including chondrogenesis. However, co-treatment with NAC did not rescue impacts of DNA damage, suggesting that the DNA damage observed in our studies was not oxidative.

Consistent with our previous findings, embryonic exposure to TBBPA disrupted several cartilage parameters, including lower jaw length (LJL), ceratohyal cartilage length (CCL), and overall cartilage area at both 0.05 and 0.5 μM of TBBPA compared with the control group. However, inter cranial distance (ICD) was markedly affected at only 0.5 μM of TBBPA compared with the control group **(Fig. 1A-D).** This suggests that the viscerocranium, which forms jaw cartilages reflected by LJL and CCL, may be more sensitive to TBBPA toxicity, whereas the neurocranium, which contributes to braincase width reflected by ICD, may be less susceptible to TBBPA-induced disruption. One possible explanation is that viscerocranium development is largely dependent on cranial neural crest cells (CNCCs), while the neurocranium has a mixed embryonic origin, arising from both CNCCs and mesoderm^33^. Therefore, mesoderm-derived components of the neurocranium may provide a buffering or compensatory effect, reducing the apparent impact of TBBPA on ICD ^33,34^. Ahr2 knockdown also rescued the TBBPA-induced reductions in LJL and CCL, indicating that these defects are mediated through Ahr signaling **(Fig. 6A and 6C)**. However, cartilage area was not restored in Ahr2 morphants exposed to TBBPA, suggesting that TBBPA may impair this cartilage parameter through additional mechanisms that are not solely dependent on canonical Ahr signaling **(Fig. 6D)**. Additionally, the angle-specific rescue observed here further suggests that Ahr signaling selectively contributes to craniofacial cartilage development. Notably, the rescue of the PQ-Meckel’s and CH-CH angles in Ahr2 morphants was consistent with the rescue of LJL and CCL, as the relative lengths and spatial orientation of LJL and CCL determine these angular measurements of PQ-Meckel’s and CH-CH angles **(Fig. 6E and 6G)**. This suggests that TBBPA disrupts distinct parameters of viscerocraniam development through an Ahr-dependent mechanism, whereas the lack of rescue in Meckel’s angle and the CH-PQ angle further indicates that additional Ahr-independent mechanisms are also involved.

Our functional enrichment analysis also revealed that TBBPA also disrupted a coherent sequence of developmental processes required for craniofacial cartilage formation **(Fig. 2B).** The enriched pathways spanned early cell fate specification and differentiation, including NCC specification, followed by biological processes essential for NCCs migration to pharyngeal arches such as epithelial to mesenchymal transition (EMT) and cell migration. These early developmental alterations were further accompanied by enrichment of downstream biological processes associated in cartilage development such as mesenchymal cell and chondrocyte differentiation, cartilage development, and embryonic viscerocranium morphogenesis. Since craniofacial cartilage develops predominantly from migratory cranial neural crest cells ^33^, disruption of this developmental trajectory provides a mechanistic link between Ahr activation, ROS production, and the observed craniofacial cartilage abnormalities. TBBPA-induced changes in E-cadherin and N-cadherin were consistent with our previous study ^19^ and these effects were rescued by Ahr2 knockdown **(Fig. 7A-C)**, further supporting an Ahr-dependent role in TBBPA-induced disruption of neural crest migration and chondrogenesis. Interestingly, Snail2-a key transcription factor for EMT, displayed a different pattern, with increased expression in DMSO-treated Ahr2 morphants but no change due to TBBPA **(Fig. 7C)**, suggesting that AHR-mediated EMT effects may be acting through other EMT transcription factors. In addition to these EMT markers, Ahr2 knockdown also mitigated effects on Sox2 and Sox10, indicating that TBBPA-induced disruption of early neural crest and chondrogenic markers is dependent on Ahr signaling. Consistent with this interpretation, previous studies have shown that Ahr expression is repressed by pluripotency factors such as Sox2 in embryonic stem cells, suggesting that precise temporal regulation of Ahr signaling is essential for normal embryonic development and cellular differentiation ^35^. Downstream during development, TBBPA also reduced the expression of chondrocyte-associated marker, Col2a1 ^36^, that was restored by Ahr2 knockdown **(Fig. 7F)**. Col2a1 is a key cartilage matrix marker expressed by developing chondrocytes ^37^. Therefore, its decreased expression and rescue by Ahr2 knockdown reinforces our phenotypic data that cartilage formation is disrupted in an Ahr-dependent manner. These collective observations are consistent with a previous study^38^ demonstrating that TCDD; a potent Ahr agonist exposure disrupts multiple stages of chondrogenesis, including mesenchymal cell recruitment, chondrocyte proliferation, differentiation, and maturation into hypertrophic chondrocytes, in a concentration-dependent manner ^38^. Finally, we sought to examine if ROS plays a role in TBBPA-induced cartilage defects, since prior studies have shown that redox homeostasis is a crucial regulator of cartilage development ^39^. However, in contrast to Ahr2 knockdown, NAC rescued only the decrease in CCL and increase in PQ-Meckel’s angle, whereas the other cartilage parameters remained disrupted **(Fig. 10C and 10E)**. Together, these findings suggest that-1) AHR signaling regulates TBBPA-induced craniofacial toxicity by disrupting both early cranial neural crest cell processes, including EMT and migration, and later chondrogenic differentiation required for normal cartilage formation and some of these effects are modulated by AHR-mediated ROS.

## 5. Conclusion

This study demonstrates that environmentally relevant TBBPA exposure disrupts craniofacial cartilage development through activation of the AHR-Cyp1a signaling axis. Ahr2 knockdown rescued TBBPA-induced Cyp1a expression, ROS production, DNA damage, chondrogenic marker alterations, and select cartilage defects, supporting an Ahr2-dependent mechanism. In contrast, the direct effect of oxidative damage on downstream phenotypes, including chondrogenesis, is prevalent, but limited to only specific morphometric and molecular parameters. Future studies should further investigate downstream mechanisms linking Ahr2 activation to impaired chondrogenesis, including the potential involvement of slincR (sox9b long intergenic noncoding RNA). Given our findings, additional work is needed to study the relative contributions of Ahr-dependent and ROS-independent pathways to TBBPA-induced cartilage developmental defects as slincR is induced by the Ahr signaling. Future studies should also determine whether short-term TBBPA exposure produces persistent effects on craniofacial cartilage development or whether these alterations can recover after the exposure is discontinued.

## 6. CRediT authorship contribution statement

Kanchaka Senarath Pathirajage: Writing – review & editing, Writing – original draft, Visualization, Validation, Methodology, Investigation, Formal analysis, Data curation, Conceptualization. Sunil Sharma: Methodology. Tyler Johnson: Investigation. Arnob Sarker: Methodology, Investigation. Subham Dasgupta: Writing – review & editing, Writing – original draft, Visualization, Validation, Supervision, Software, Resources, Project administration, Methodology, Investigation, Funding acquisition, Formal analysis, Data curation, Conceptualization.

## 9. Acknowledgements

We acknowledge Mr. Scott Horman of Clemson’s Aquatic Animal Research Lab for their role in fish husbandry.

